# Family Conflict and Parent-Child Similarity in Affective Valuation

**DOI:** 10.64898/2026.06.24.734340

**Authors:** Tae-Ho Lee, Ya-Yun Chen, Qingyi Li, Beiming Yang, Zexi Zhou, Yang Qu

## Abstract

How children come to evaluate social-affective cues is shaped within the family, yet the neural expression of this process and its dependence on family relationships remain unclear. We tested whether parent-child similarity in the neural coding of affective judgment varies with family environment, and whether it relates to youth affective distress. Twenty-five parent-child dyads (youth, mean age 11.8; parents mean age 42 years) judged faces morphed along an angry-to-happy continuum as positive or negative during fMRI. For each participant, we defined an evaluative choice axis distinguishing faces judged positive from negative, independent of expression intensity. Using searchlight-based parent-child cross-decoding, we tested whether one dyad member’s evaluative coding predicted the other member’s judgments, indexing shared evaluative coding rather than shared sensitivity to expression intensity. Inference focused a priori on medial prefrontal cortex. There was no reliable average parent-child neural similarity across the sample. Instead, higher family conflict was associated with lower parent-child neural similarity in ventromedial prefrontal cortex (vmPFC). Demonstrating specificity to negative relational strain, this effect was not observed for complementary dimensions of family cohesion or identity. Moreover, the effect was specific to true dyads rather than random pairings and to vmPFC rather than a face-selective network or other medial prefrontal regions. Lower vmPFC similarity showed a preliminary association with higher youth affective distress. These findings indicate that affective valuation, rather than sensory encoding, may be a representational level at which perceived family conflict is reflected in parent-child neural similarity.

## Family Conflict and Parent-Child Similarity in Affective Valuation

Social-affective perception requires people to interpret cues whose meaning is often graded, context-dependent, and shaped by interpersonal experience. A tone of voice, facial expression, gaze pattern, body posture, or pause in conversation can carry affective meaning, but the same cues can be interpreted differently across perceivers (Aviezer et al., 2008; Ko et al., 2011; Lee et al., 2012; Lim & Pessoa, 2008). This is because perception is not a passive registration of sensory input, but an active interpretation of sensory evidence shaped by the perceiver’s prior experience, expectations, and values (Barrett et al., 2011; Bruner & Goodman, 1947; De Lange et al., 2018). Such perceiver-dependent differences may be most apparent when cues are subtle or mixed (Neta & Whalen, 2010), but they are not limited to ambiguous cues. This perceiver-dependence is not only behavioral. When people view the same face and arrive at different judgments, those judgments are accompanied by distinct neural responses that are dissociable from the activity driven by the stimulus itself (Thielscher & Pessoa, 2007). In other words, how a face is evaluated, not only what is physically present, is reflected in the brain. Across social-affective perception more broadly, affective meaning does not arise from sensory features alone. It depends on which cues are selected, weighted, integrated, and interpreted as emotionally meaningful (Barrett & Bar, 2009).

## The Family Context of Neural Affective Valuation

Family environments provide a primary relational setting in which children learn how these social-affective cues should be evaluated. Parent-child relationships involve repeated exposure not only to the same social situations, but also to caregivers’ ways of evaluating social-affective cues, responding to uncertain affective information, and assigning affective value to social signals (Ehli, 2020; Eisenberg et al., 1998; Morris et al., 2007; Mumme et al., 1996; Sorce et al., 1985). Over time, these relational experiences typically contribute to similarity between parents and children in how social-affective information is evaluated and interpreted (Grusec & Goodnow, 1994). Children tend to internalize their parents’ evaluative standards, particularly earlier in development, and this input shapes their own valuation (Lim et al., 2016). Through this internalization, children’s own evaluative criteria come to resemble those of their parents (Qu et al., 2023; Zhou et al., 2026), and the resulting parent-child similarity has also been documented at the neural level (Chen, 2025; Clinchard et al., 2024; Lee, Miernicki, et al., 2017a, 2017b; Lee et al., 2018; Zhou et al., 2023)

For affective evaluation specifically, the relevant neural system supporting this valuation is the medial prefrontal cortex (mPFC). Within this system, the ventromedial prefrontal cortex (vmPFC) is one of the most consistent sites of subjective-value representation across domains (Amodio & Frith, 2006); (Bartra et al., 2013; Clithero & Rangel, 2014). Because the affective judgments of interest concern whether a cue is evaluated as positive or negative, vmPFC value representation is the primary target. If the criteria by which affective cues are evaluated are shaped within the family, parent-child similarity—or divergence—in this evaluation should be expressed in vmPFC patterns, rather than in perceptual regions that merely encode visual features.

## Family Conflict and Neural Similarity in Affective Evaluation

Importantly, children’s internalization of evaluative standards is not a uniform process; whether parental standards are taken up depends heavily on the child’s acceptance of them, which varies with the quality of the parent-child relationship (Darling & Steinberg, 1993; Davies & Cummings, 1994; Grych & Fincham, 1990). Family conflict, which is characterized by frequent negative interactions, disagreements, and relational strain (Olson et al., 1979; Ruiz et al., 1998; Tyler & Degoey, 1995), may disrupt this emotional socialization process. In high-conflict environments, youth may actively reject parental evaluative standards, or caregivers’ emotional cues may be perceived as too hostile or inconsistent to serve as reliable guides. Consequently, youth’s developing architecture for assigning affective value may diverge from their parents’. For example, recent work has linked negative family emotional environment to reduced medial prefrontal child-parent synchrony (Su et al., 2024), suggesting that high conflict acts as a developmental barrier to the formation of shared affective valuation.

In addition, it is equally plausible that an existing divergence in neural evaluative coding actively contributes to family conflict. Because affective perception is a perceiver-dependent interpretation, a lack of shared evaluative coding means that parents and youth will frequently decode the same graded, everyday social cues differently. A parent might perceive a neutral or mixed expression as benign, while the youth perceives it as threatening, which is a negativity bias that is particularly pronounced for ambiguous expressions during childhood and adolescence (Neta & Whalen, 2010) According to social information-processing theory, the tendency to misinterpret ambiguous social cues toward hostile or negative intent is heavily influenced by prior interpersonal experiences, and this hostile attribution bias acts as a primary catalyst for aggressive behavior and interpersonal friction (Coe et al., 2020; Yaros et al., 2016). When parents and children lack a shared neural choice axis for what constitutes a positive versus negative cue, their continuous misalignments in emotional reactions likely precipitate arguments and misunderstandings (Crick & Dodge, 1994; Dodge et al., 2015). Together, these two theoretical pathways suggest that family conflict and shared neural evaluative coding could be inversely related. Establishing whether an association exists between negative family environment and divergent affective valuation is a critical step in understanding adolescents’ affective functioning.

## Parent-Child Neural Similarity and Youth Affective Distress

Parent-child neural similarity in affective valuation may also have development significance and be relevant for youth mental health (Clinchard et al., 2024; Kim-Spoon et al., 2024). Prior research has demonstrated that parent-child neural similarity during emotionally evocative tasks is linked to youth emotional adjustment (Chen et al., 2025; Clinchard et al., 2024; Lee, Miernicki, et al., 2017a, 2017b; Lee et al., 2018; Zhou et al., 2023). Medial prefrontal systems continue to mature across childhood and adolescence, a period in which affective evaluation is intensely shaped by emotion socialization (Kilford et al., 2016). Because vmPFC supports subjective valuation, similarity between parents and youth in vmPFC evaluative patterns may reflect a shared framework for assigning positive versus negative meaning to social cues. Such shared coding may support mutual understanding and provide youth with more consistent relational feedback about emotionally meaningful information. In contrast, lower parent-child vmPFC similarity may indicate divergence in how social-affective cues are valued, potentially contributing to friction, uncertainty, and negative interpretations, that consequently exact a toll on adolescent mental health. Understanding whether shared or divergent evaluative coding relates to youth affective distress provides critical insight into how the dyadic family environment influences adolescents’ broader socioemotional well-being.

## The Present Study

In the present study, we tested whether parents and children use similar neural patterns when assigning affective meaning to social-affective cues, and whether this dyadic neural similarity varies as a function of family environment and is associated with youth affective adjustment. Twenty-five parent-child dyads with youth from middle childhood through adolescence completed an fMRI facial-affect perception task in which they judged faces morphed along an angry-to-happy continuum as positive or negative. Rather than assessing facial-expression discrimination accuracy, we aimed to capture neural processes involved in evaluating affective meaning. For each participant, we defined an evaluative choice axis distinguishing faces judged positive from negative, independent of the physical intensity of the expression. We then used searchlight-based parent-child cross-decoding to test whether one dyad member’s evaluative coding predicted the other member’s judgments, providing an index of shared evaluative mapping.

Inference focused a priori on the mPFC, given its central role in affective evaluation and social meaning assignment. Based on the theoretical link between relational strain and divergent affective perception, we primarily hypothesized that greater family conflict would be associated with lower parent-child neural similarity in the vmPFC. To evaluate whether this dyadic neural divergence is specifically sensitive to the negative relational strain of conflict, we also tested family cohesion and family identity as separately examined, complementary predictors (Olson et al., 1979; Ruiz et al., 1998; Tyler & Degoey, 1995). This analytical approach allowed us to determine if neural similarity is specifically responsive to conflict, rather than broadly associated with positive family environment dimensions like perceived closeness, support, and the youth’s sense of belonging. A face-selective perceptual network also served as a specificity control to test whether any family- environment association was specific to evaluative regions rather than sensory-perceptual processing of facial features (Julian et al., 2012). Finally, to evaluate the developmental relevance of these dyadic neural patterns, we tested whether vmPFC neural similarity was associated with a composite measure of youth affective distress. We hypothesized that greater parent-child vmPFC neural similarity would be linked to lower levels of youth affective distress.

## Methods

### Participants

Twenty-six parent-child dyads completed the face emotion perception task. One dyad was excluded because one dyad member did not complete the task, stopping at trial 47 of 196. The final analytic sample included 25 parent-child dyads, with children and adolescents aged 8-17 years, from middle childhood through adolescence, hereafter referred to as youth (youth: M_age_ = 11.84 years, SD = 2.56, range = 8-17, 40% female; parents defined as the self-identified primary caregiver: M_age_ = 41.96 years, SD = 5.68, range = 33-54, 52% female; **Fig1.A**).

**Fig 1.**
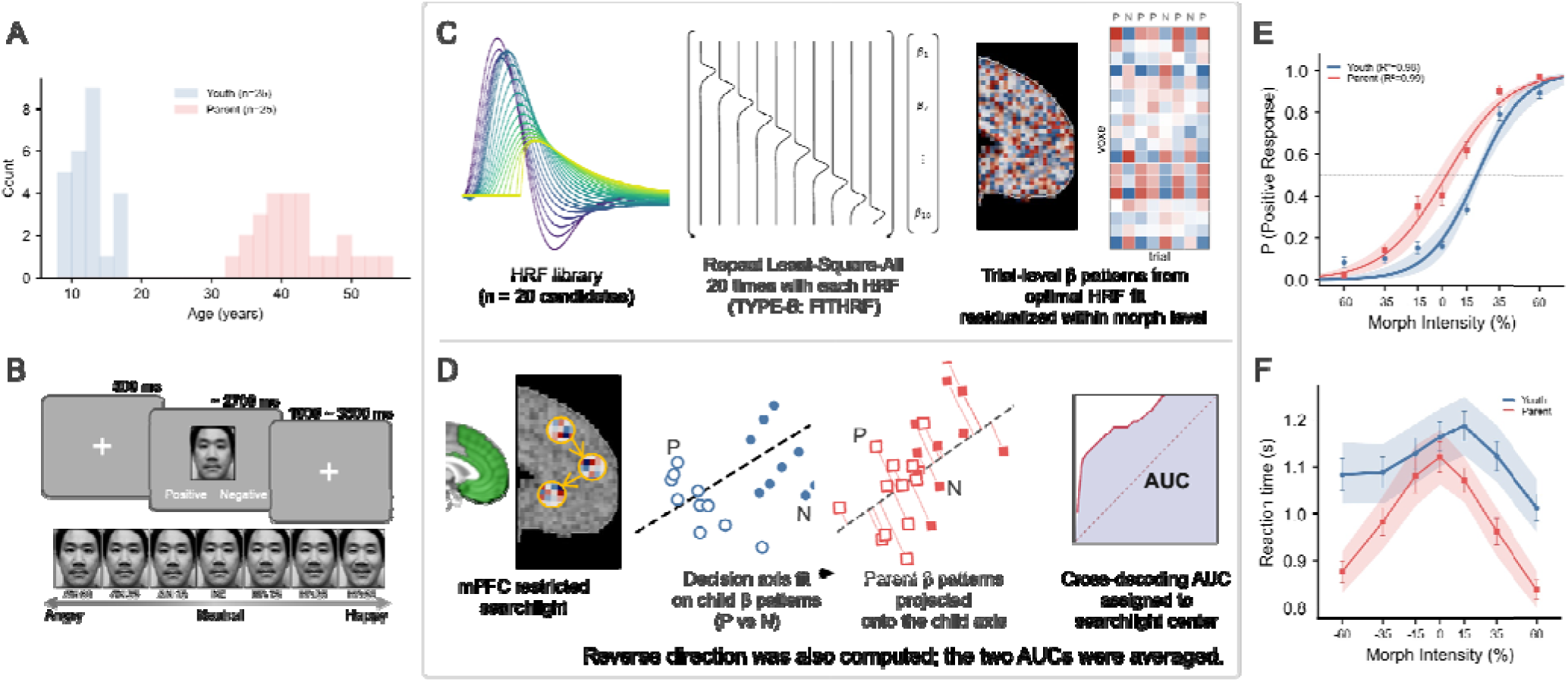
Task design, cross-decoding analysis pipeline, and behavioral performance. (A) Age distribution of youth and parent participants included in the final analytic sample. (B) Facial-affect judgment task. Participants viewed faces morphed along an angry-to-happy continuum and categorized each face as positive or negative. (C) Single-trial neural pattern estimation. Trial-level β patterns were estimated using GLMsingle (TYPE-B, FITHRF), in which candidate HRFs were evaluated voxel by voxel and the best-fitting HRF for each voxel’s BOLD time series was used to obtain trial-level β estimates. P = trials judged positive, N = trials judged negative. (D) mPFC searchlight cross-decoding. Searchlight analyses were performed within the a priori mPFC mask. Within each searchlight, patterns were mean-centered within each morph-intensity level before defining positive-versus-negative evaluative axes. P and N indicate trials judged positive and negative, respectively. The schematic illustrates child-axis to parent-trial decoding: the child’s evaluative axis was applied to the parent’s trial-level β patterns, and AUC was computed from the ranked projection scores before being assigned to the searchlight center. The reverse parent-axis to child-trial direction was also computed, and the two directional AUC values were averaged to form the dyad-level parent-child neural similarity map. (E) Group-mean psychometric functions, with 95% confidence bands for youth and parents across the angry-to-happy morph continuum. Points indicate mean positive-response rates at each morph intensity, and fitted curves show logistic model estimates. (F) Mean reaction time across morph intensity. Reaction times were longest near the most ambiguous morphs and decreased toward the less ambiguous endpoints in both groups. Note. Photographs shown in the task schematic are of the authors and are reproduced with their permission.

### Task and Stimuli

Participants completed a parametric forced-choice emotion-categorization task using faces morphed along an angry-to-happy continuum (**Fig1.B**). On each trial, participants judged each face as positive or negative. To vary emotional intensity parametrically, angry and happy faces were morphed with neutral faces in 1% increments by linearly interpolating facial landmark positions and pixel intensities between photographs of the same individual displaying angry, neutral, and happy expressions using Delaunay triangulation-based morphing. From these morph continua, we selected three intensity levels for each emotion category (15%, 35%, and 60%) and additionally included a neutral face. The source set used to generate the stimuli consisted of 14 face identities, including 7 female and 7 male identities, drawn from established facial expression datasets (Lundqvist et al., 1998; Tottenham et al., 2009).

Each face was presented in grayscale within a rectangular frame (3.0° × 3.3°) that excluded most hair and other nonfacial contours. This yielded 98 unique face stimuli reflecting 14 identities at seven morph levels, and each stimulus was presented twice across the task, resulting in 28 trials per intensity level and 196 trials per participant. Participants indicated by button press whether each face appeared positive or negative. On each trial, a fixation cross was presented for 500 ms, followed by a morphed face displayed for up to 2,700 ms, with the trial ending upon response. Intertrial intervals were jittered between 1 and 3.5 s.

### Questionnaire Measures

#### Family conflict

Family conflict was measured using the Parent-Adolescent Conflict Scale (PACS; Ruiz et al., 1998). On a 5-point Likert scale ranging from 1 (almost never) to 5 (almost always), youth reported on the frequency of conflict between them (10 items, e.g., “you and your parent had a serious argument or fight” and “you and your parent yelled or raised voices at each other”). The mean was taken across all items, with higher scores indicating more frequent perceived family conflict (αs□=L0.82, *M* = 2.04, *SD* = 0.66, and range = 1.10–3.60).

#### Complementary family environment dimensions

Family cohesion was measured using 10 items in the cohesion subscale of the Family Adaptation and Cohesion Evaluation Scales II inventory (Olson et al., 1979). On a 5-point Likert scale ranging from 1 (almost never) to 5 (almost always), youth reported on their relationship closeness with parents (e.g., “My mother/father and I are supportive of each other during difficult times”). The scores were averaged across all items, with higher scores indicating greater family cohesion (α□=□0.73, *M* = 3.90, *SD* = 0.78, and range = 1.70–4.80). Youth also reported on family identity using a measure assessing perceived belonging, value, and recognition within the family (8 items; e.g., “I feel that I belong in my family”) on a 5-point Likert scale ranging from 1 (Strongly Disagree) to 5 (Strongly Agress)(Tyler & Degoey, 1995). The mean was taken across all items, with higher scores indicating stronger family identification (αs□=□0.95, *M* = 3.86, *SD* = 1.21, and range = 1.00–5.00).

#### Youth’s affective distress

To capture affective distress in youth on a single dimension, we combined five youth-report measures into a affective distress composite: self-reported depression (Barkmann et al., 2008), state- and trait-anxiety (Spielberger et al., 1973), negative affect (10 items; e.g., “distressed”, “upset” ; Laurent et al., 1999) and perceived stress (Cohen et al., 1983). Each measure was z-scored across the 25 children, and the composite was computed as the unweighted mean of the five z-scores. Descriptive statistics for the five affective-distress components were as follows: depression, *M* = 14.92, *SD* = 12.47, range = 0-57; state anxiety, *M* = 33.08, *SD* = 9.46, range = 22-56; trait anxiety, *M* = 38.32, *SD* = 8.15, range = 26-60; negative affect, *M* = 30.20, *SD* = 13.32, range = 15-69; perceived stress, *M* = 17.60, *SD* = 7.86, range = 2-39. The youth affective-distress composite, computed as the mean of the z-scored components, had *M* = 0.00, *SD* = 0.84, range = −1.23 to 2.44, and showed good internal consistency with Cronbach’s α = .89.

### fMRI Acquisition and Preprocessing

All MRI data were acquired on a Siemens 3T PRISMA with a 64-channel matrix head coil. High-resolution T1w (TR=2.5 s; TE=2.06 ms; FA=8°; 1 mm isotropic voxel) and T2w (TR=3.2 s; TE=563 ms; FA=120°; 1 mm isotropic voxel) anatomic images were acquired for tissue segmentation (GM, WM, and CSF mask) and normalization. Functional images were acquired with gradient-echo echo-planar T2*-weighted imaging sequence (TR=2s; TE=25 ms; FA=90°; 2.5 x 2.5 mm resolution; 37 interleaved 3.0 mm slices with 0.3 mm gap).

Preprocessing was performed using the FMRIB Software Library (FSL; Jenkinson et al., 2012), ICA-AROMA toolbox (Pruim et al., 2015), and ANTs library (Avants et al., 2009). The first two volumes were removed, and images were then high-pass filtered with a 128-s cutoff, motion-corrected, grand-mean intensity-normalized, denoised using ICA-AROMA After AROMA denoising (corrected mean framewise displacement: Children = 0.039 ± 0.019 mm; parents = 0.027 ± 0.007 mm); computed linear/non-linear registration matrices based on the standard 2-mm MNI brain template. Spatial smoothing was not applied because the primary analyses relied on local multivoxel activation patterns.

### Trail-by-Trial Neural Pattern Estimation

Single-trial neural patterns were estimated separately for each participant using GLMsingle (Prince et al., 2022), which combines a voxel-wise HRF library with single-trial GLM estimation (**Fig1.C**). Rather than estimating one average response per stimulus or response condition, each trial was entered as a separate regressor in a single participant-level design matrix (LSA-style; up to 196 trial-specific regressors). This yielded one β estimate for each trial and voxel. The model was fit to unsmoothed native-space EPI data, with six rigid-body motion parameters and framewise outlier regressors included as nuisance covariates.

GLMsingle’s library of 20 candidate double-gamma HRFs covers the empirical range of voxel-wise hemodynamic shapes (Prince et al., 2022). For each voxel, the full single-trial GLM was fit separately with each candidate HRF, and the HRF that best fit the BOLD time series, defined by the highest variance explained, was retained. The β estimates for that voxel were then taken from the corresponding fit. Thus, HRF shape was voxel-specific but held constant across trials within a voxel, allowing local hemodynamic variation to be modeled without treating HRF differences as trial-by-trial fluctuations. We used these TYPE-B FITHRF β estimates as the trial-level neural patterns.

Trial-wise β images were aligned with each participant’s trial labels, including morph intensity and positive versus negative response, and served as the input to the response-defined parent-child cross-decoding analysis. Estimation was performed in native EPI space to preserve fine-grained multivoxel structure that may be attenuated by resampling. The resulting β images were then warped to the 2-mm MNI152 template using each participant’s ANTs native-to-MNI transform chain for group-level searchlight and ROI analyses. Additional details of the single-trial β estimation and parent-child cross-decoding workflow are provided in **Supplementary Fig S1.**

### Parent-Child Neural Similarity From Choice-Axis Cross-Decoding

The analysis was designed to capture how a participant evaluated a face as positive or negative, rather than sensitivity to the physical intensity of the expression. For each participant, trial-level beta patterns were first mean-centered within each morph-intensity condition. Specifically, for each of the seven morph levels, we subtracted that participant’s average neural pattern for that morph level from each trial at the same level. This reduced stimulus-driven pattern structure associated with facial-expression intensity and allowed the analysis to focus on trial-to-trial variation related to whether the face was judged as positive or negative. This procedure follows the broader logic of controlling nuisance-related pattern structure in multivariate decoding analyses, where otherwise decodable information can reflect confounded stimulus or task properties rather than the process of interest (Snoek et al., 2019; Todd et al., 2013). Within each 10-mm-radius searchlight sphere, we then defined a participant-specific evaluative choice axis as the voxel-wise difference between the mean condition-centered pattern for faces judged as positive and the mean condition-centered pattern for faces judged as negative (**Fig1.D**), following the logic of using choice-related variation in neural responses to identify how a perceptual decision is made when the stimulus is held comparable (Thielscher & Pessoa, 2007), here extended to test whether this evaluative coding is shared across parent-child dyads. Whereas that work related single-region response amplitude to perceptual choice, we defined a multivariate pattern axis and tested its generalization across individuals. Because the morph-intensity mean had been removed within each level, two trials at the same intensity could still differ in whether the face was judged positive or negative, and the evaluative choice axis captured the neural pattern accompanying this difference in judgment rather than the physical expression itself. The dyadic question was then whether the axis distinguishing one member’s positive from negative judgments also distinguished the other member’s judgments, indicating that the two organized affective meaning in similar neural terms.

Parent-child neural similarity was operationalized as cross-decoding-based similarity. For each dyad, child-axis to parent-trial area-under-curve (AUC) was computed by applying the child’s evaluative choice axis to the parent’s condition-centered trial patterns and evaluating whether the resulting projection scores separated the parent’s positive- from negative-response trials. Parent-axis to child-trial AUC was computed analogously by applying the parent’s evaluative choice axis to the child’s condition-centered trial patterns and evaluating whether the resulting projection scores separated the child’s positive- from negative-response trials. The two directional AUC maps were averaged to form the primary dyad-level neural similarity map, with AUC = 0.5 indicating chance-level cross-decoding.

Group inference tested whether these dyad-level neural similarity maps varied as a function of family environment, using the permutation-based cluster procedure described above (Winkler et al., 2014). Small-volume correction was applied within an a priori Brainnetome medial prefrontal mask, defined as the voxel-wise union of bilateral anterior medial prefrontal cortex (amPFC; 1,999 voxels), dorsal medial prefrontal cortex (dmPFC; 4,648 voxels), and ventral medial prefrontal cortex (vmPFC; 5,280 voxels), for a total mask size of 11,927 voxels (Fan et al., 2016).. For each covariate model, covariate-positive and covariate-negative contrasts were tested.

### Statistical Analysis and Reporting

Unless otherwise specified, correlations are Pearson coefficients and all tests are two-tailed, with significance evaluated at p < .05. All reported 95% confidence intervals were obtained by bootstrap resampling (10,000 resamples) and are referred to as 95% CIs throughout. Permutation-based inference was used for group-level cluster analyses (5,000 permutations; FSL randomise, cluster-defining threshold z > 3.1, small-volume cluster-FWE p < .05 within the a priori medial-prefrontal mask) and for the random-pair specificity analysis (10,000 derangements excluding all true dyads). The three family-environment subscales (conflict, cohesion, identity) were examined as a priori, separately tested dimensions; no additional correction was applied across the three subscales. Robustness of the primary association was assessed with the random-pair derangement analysis and with leave-one-dyad-out analyses. The youth affective distress composite was computed as the unweighted mean of z-scored measures of depression (CES-DC), state anxiety (STAI-S), trait anxiety (STAI-T), negative affect (PANAS-N), and perceived stress (PSS); each component was z-scored within the n = 25 youth distribution before averaging (Cronbach’s α = .89).

## Results

### Behavioral Characteristics

Because the primary goal of the task was to characterize parent-child neural similarity during positive versus negative affective judgments, rather than individual differences in expression-intensity discrimination, behavioral analyses were used to confirm that both children and parents performed the task systematically across the morph continuum. For each participant, trial-level binary responses (positive vs. negative) across the seven signed morph levels (AN60 = −60, AN35 = −35, AN15 = −15, NE = 0, HA15 = +15, HA35 = +35, HA60 = +60; signed morph units) were fit with a logistic function, P(happy | x) = 1 / (1 + exp(−k(x − PSE))). PSE indexed the morph level at which a participant was equally likely to categorize a face as positive or negative, and k indexed sensitivity to expression-related visual feature changes along the morph continuum.

Children showed a stronger negativity-biased response pattern than parents (**Fig.1E**). Their PSE was shifted further toward the happy end of the morph continuum, indicating that children used a more negative response criterion when categorizing mixed facial expressions as positive versus negative (PSE: child M = 20.1 morph units, SD = 15.1; parent M = 1.5, SD = 17.5; paired t(24) = 4.35, p < .001; 95% CI for child-parent difference [10.3, 26.8]). This direction of bias is consistent with previously reported developmental patterns of negative interpretation bias (Tottenham et al., 2013). Psychometric steepness did not differ between children and parents (k: child M = 0.089, SD = 0.043; parent M = 0.075, SD = 0.033; paired t(24) = 1.25, p = .22), indicating no evidence that children and parents differed in sensitivity to expression-related visual feature changes. Mean reaction time was longer for children than for parents (child M = 1.17 s, SE = 0.04; parent M = 1.01 s, SE = 0.02; paired t(24) = 3.66, p = .001; 95% CI [0.05, 0.19]), with reaction times longest at the most ambiguous morphs and decreasing toward the unambiguous extremes in both groups (**Fig1.F**).

Together, these behavioral results indicate that both youth and parents used the morph continuum systematically, supporting the use of the task to examine parent-child neural similarity during affective judgment.

### Parent-Child Neural Similarity and Family Conflict

The main parent-child cross-decoding searchlight analysis did not yield any surviving clusters after small-volume correction within the a priori medial prefrontal mask, indicating no reliable average dyadic neural similarity effect. Family-environment covariate analyses were then used to test whether variability in cross-decoding-based neural similarity was associated with children’s perceived family conflict. This analysis yielded a surviving cluster in vmPFC (84 voxels; peak MNI = 6, 46, −16; peak t = 5.24; cluster-level FWE p < .05; **Fig.2A**). The effect was negative, indicating lower parent-child neural similarity in dyads with higher family conflict.

**Fig 2.**
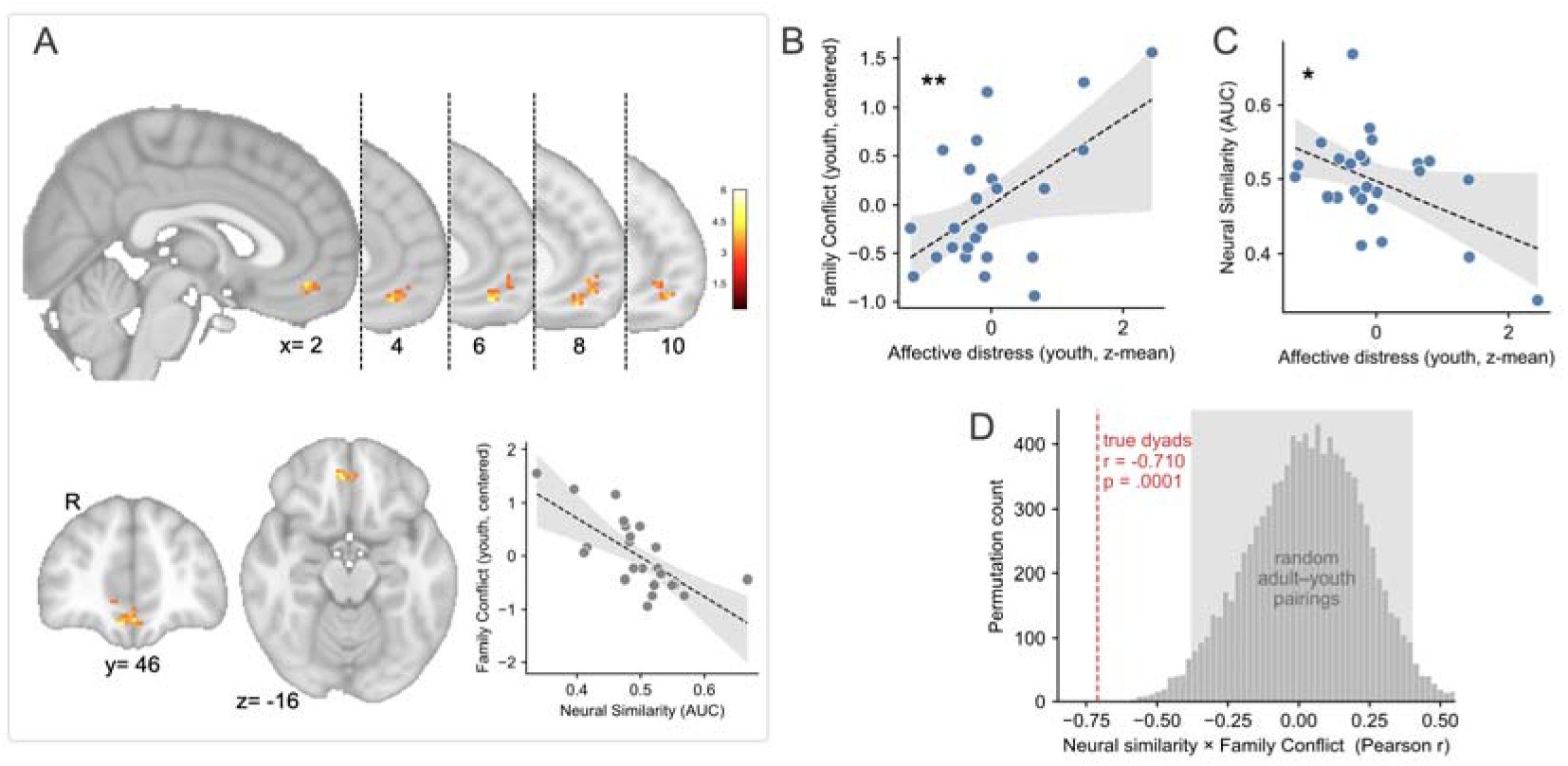
Parent-child neural similarity in vmPFC decreases with family conflict. (A) Small-volume permutation-corrected vmPFC cluster within the a priori medial prefrontal mask where parent-child similarity decreased as family conflict increased. The adjacent scatterplot, showing cluster-mean neural neural similarity against conflict, is for visualization only; statistical inference for the vmPFC cluster was based on the searchlight analysis, not on the plotted correlation. (B) Association between family conflict and youth affective distress. (C) Association between parent-child neural similarity and youth affective distress. This association reached a nominal threshold but was not stable under resampling and should be interpreted as preliminary (see text). (D) Random-pair analysis showing that the family conflict association with neural similarity was observed for true parent-child dyads but not for random adult-youth pairings (derangements excluding all true dyads). \**p* < .05, \*\**p* < .01.

To examine the specificity of this association, we conducted parallel analyses for other family-environment variables. No surviving clusters were observed for family cohesion or family identity, suggesting that the vmPFC neural similarity effect was specific to family conflict rather than reflecting family environment more broadly.

To characterize whether the neural similarity-family conflict association in vmPFC was driven by one decoding direction, we decomposed the direction-averaged neural similarity index into child-axis to parent-trial decoding and parent-axis to child-trial decoding. The association with family conflict was present in both directions, indicating that the effect was not attributable to one decoding direction alone. Full directional results are reported in **Supplementary Table S1.**

### Parent-Child Neural Similarity and Youth’s Affective Distress

We next examined associations among youth affective distress, family environment, and parent-child neural similarity. Youth affective distress was positively associated with family conflict (r = 0.559, p = .004; 95% CI [0.04, 0.82]; **Fig.2B**), but was not significantly associated with family cohesion (r = −0.152, p = .47) or family identity (r = −0.044, p = .83). We then tested whether parent-child neural similarity was associated with youth affective distress. Mean cross-decoding-based neural similarity was extracted from the vmPFC cluster identified in the preceding family-environment covariate analysis and correlated with the youth affective distress composite (defined above). The Pearson correlation reached the conventional p < .05 threshold (r = −0.480, p = .015; **Fig.2C**), but the bootstrap 95% confidence interval included zero ([−0.75, 0.01]), indicating that the composite-level association was not stable under resampling. Thus, the association between lower vmPFC parent-child neural similarity and higher youth affective distress should be interpreted as preliminary. Constituent-scale associations are reported in **Supplementary Table S2.**

### Specificity and Robustness

#### Random-pair analysis

We conducted a random-pair analysis to evaluate whether the association depended on true parent-child pairing rather than arbitrary adult-youth matching. This analysis tested whether the observed association reflected a dyad-specific relation between family conflict and neural similarity, or whether a similar association would arise when youth were paired with unrelated adults from the same sample. The observed true-dyad association (r = −0.710) fell outside the null distribution generated from 10,000 derangements that excluded all true dyads (null mean r = 0.028, 95% interval [−0.380, 0.402]; permutation p = .0001; **Fig.2D**). Mean neural similarity was near chance for both true dyads (M = 0.497) and random pairs (M = 0.501), indicating that the result was not explained by higher average cross-decoding in true dyads. Thus, the family conflict association was observed for genuine parent-child dyads but not for random adult-youth pairs, supporting the interpretation that the vmPFC effect reflects dyad-specific variation in parent-child evaluative-pattern similarity rather than a generic adult-youth cross-decoding pattern.

#### Face-selective network control

To evaluate whether the neural similarity-family conflict association in vmPFC could be explained by sensory-perceptual similarity for faces, we repeated the cross-decoding analysis within a face-selective network parcellation (Julian et al., 2012), covering bilateral face-selective regions in occipitotemporal cortex, including FFA, OFA, and posterior STS (5,900 voxels in 2-mm MNI space). No cluster survived FWE correction in either direction. Mean face-network neural similarity was at chance (M = 0.504; one-sample t = 1.42, p = .17) and was not associated with any of the three family-environment subscales (family conflict r = −0.193, p = .36; family cohesion r = -0.214, p = .30; family identity r = 0.125, p = .55). Thus, parent-child similarity in face-selective sensory-perceptual regions neither exceeded chance nor tracked family environment, indicating that the vmPFC neural similarity-family conflict association did not reflect shared sensory-perceptual processing of facial features.

#### Leave-one-dyad-out robustness

As an additional robustness check, we conducted a leave-one-dyad-out analysis to test whether the association between vmPFC parent-child neural similarity and family conflict was driven by any individual dyad. The association remained negative and significant in every leave-one-dyad-out iteration (r range = −0.769 to −0.612), indicating that the result was not driven by a single dyad. In contrast, the association between neural similarity and youth affective distress was less stable, remaining negative across iterations but losing significance in 1 of 25 iterations. These results support the robustness of the significant association between neural similarity and family conflict and are consistent with the preliminary interpretation of the youth affective distress association. Full leave-one-dyad-out results are reported in **Supplementary Table S3.**

#### Dyad-level behavioral covariate controls

We also tested whether the vmPFC- family conflict association was accounted for by dyad-level task-behavioral characteristics. The association remained significant after controlling separately for dyad mean reaction time, absolute parent-youth PSE difference, absolute parent-youth psychometric steepness difference, and absolute parent-youth reaction-time difference (partial r range = −0.688 to −0.709, all ps < .001). Thus, the vmPFC- family conflict association was not accounted for by dyad-level response speed or parent-youth differences in behavioral task performance. Full covariate-control results are reported in **Supplementary Table S4.**

## Discussion

The present study tested whether parent-child neural similarity during affective judgment varied as a function of family conflict and examined its potential developmental relevance for youth mental health. The main finding was that greater family conflict was associated with lower parent-child neural similarity in vmPFC during positive versus negative judgments of emotional faces. This association was not observed for family cohesion or family identity, and the main cross-decoding analysis did not show a reliable average parent-child neural similarity effect across the full sample. Thus, the result is not that parent-child dyads uniformly shared an evaluative neural code. Rather, parent-child neural similarity varied with youth-perceived family conflict. Additionally, lower vmPFC similarity showed a significant but less resampling-robust association with greater youth affective distress, providing tentative evidence that divergence in parent-child affective valuation may have implications for youth socioemotional functioning.

### Context and Localization of Evaluative Neural Similarity

This finding extends prior work showing that parent-child neural similarity is not a fixed property of dyads, but varies with relational and emotional context (Zhou et al., 2026). Previous fMRI studies have shown that parent-child neural similarity is stronger in families with higher connectedness during stress-related processing and that parent-child neural similarity during emotionally evocative movie viewing is linked to youth emotional adjustment, particularly in more cohesive family contexts (Lee, Qu, et al., 2017; Lee et al., 2018; Zhou et al., 2023). The present study adds to this literature by focusing on a more specific representational question: instead of measuring global similarity in responses to the same stimulus stream, we asked whether one dyad member’s neural pattern for positive versus negative affective judgment generalized to the other dyad member’s judgments. This approach targets shared evaluative mapping, not simply shared exposure to the same faces.

The localization of the effect to vmPFC helps constrain its interpretation. The cross-decoding measure was designed to reduce stimulus-driven pattern structure tied to morph intensity and to focus on trial-to-trial variation associated with positive versus negative judgment. The family-conflict association was observed in vmPFC, a region centrally involved in valuation and affective meaning assignment, but was not detected in a face-selective network. This pattern is more consistent with shared higher-level affective valuation than with shared sensory-perceptual processing of facial features. Furthermore, the effect localized to the vmPFC rather than dorsal or anterior mPFC regions, suggesting that perceived family conflict is reflected more in shared value-based evaluation than in shared social-cognitive or self-referential processing (Amodio & Frith, 2006; Bartra et al., 2013).

### The Link between Family Conflict and Evaluative Divergence

The specificity of this neural divergence to family conflict is theoretically informative. Family cohesion and family identity capture perceived connectedness, support, belonging, and family-based self-definition (Olson et al., 1979; Tyler & Degoey, 1995), whereas family conflict captures the frequency of negative parent-youth interactions, including disagreement and conflict behaviors (Ruiz et al., 1998). The absence of cohesion or identity effects suggests that shared neural valuation is uniquely sensitive to negative relational strain, rather than simply reflecting a general lack of positive closeness. Positive family environment does not necessarily imply identical evaluative coding; authoritative and supportive family environments can combine warmth with youth autonomy, allowing for normative evaluative divergence without relational damage (Bülow et al., 2022; Darling & Steinberg, 1993 ; Teuber et al., 2022).

Instead, the vmPFC finding firmly aligns with the theoretical framework of family conflict being both a potential disruptor and a potential consequence. On one hand, high family conflict may act as a developmental barrier, disrupting the normative emotional socialization processes required for a youth to internalize their caregiver’s vmPFC choice axis (Su et al., 2024). On the other hand, a lack of shared vmPFC evaluative coding may actively precipitate everyday relational friction. Because affective perception requires interpretation (Aviezer et al., 2008; Ko et al., 2011; Lee et al., 2012; Lim & Pessoa, 2008), when parents and youth use divergent neural choice axes to evaluate social-affective cues, they are highly prone to misinterpreting each other’s emotional reactions. These everyday misalignments in affective meaning assignment, as well as the potential for youth to attribute hostile intent to ambiguous cues, draw on processes that broader work has linked to threat-related cue interpretation and can actively fuel further conflict (Cisler & Koster, 2010; Fu & Pérez-Edgar, 2019). This interpretation is also supported by the behavioral data: although youth showed a more negative response criterion than parents, the neural similarity-family conflict association in the vmPFC remained significant even after controlling for these child-parent differences in point of subjective equality. Therefore, at the neural level, family conflict is tightly linked to how similarly parents and youth organize the evaluative meaning of mixed social-affective cues.

### Implications for Youth Affective Distress

The friction caused by divergent affective valuation, and the resulting cycle of interpersonal conflict, may exact a toll on adolescent mental health. Child affective distress was positively associated with family conflict, but not with family cohesion or family identity. Lower parent-child neural similarity also showed a possible association with higher child affective distress. However, although the Pearson correlation reached the conventional threshold, the bootstrap confidence interval included zero, indicating that the composite-level association was not stable under resampling. Leave-one-dyad-out analyses showed the same pattern. Thus, these findings should be treated as preliminary evidence that the dyadic neural friction associated with family conflict may be relevant to youth affective adjustment, rather than as evidence that vmPFC neural similarity robustly explains child affective distress. Disentangling whether neural similarity mediates the link between conflict and distress, whether they share a common source, or whether they are related in another way will require larger, longitudinal samples.

### Methodological Contributions and Specificity

Several analyses addressed simpler alternative explanations. First, the association was present for true parent-child dyads but not for random adult-child pairings. Parent-child neural similarity can reflect multiple sources, including shared genetics, long-term relational history, and shared environment (Zhou et al., 2026). The present design cannot separate these sources directly. However, the random-pair analysis indicates that the family conflict association was not reducible to generic adult-child similarity. Second, the association was not detected in face-selective perceptual regions, which reduces concern that the result reflects shared visual feature processing. Third, the association remained after adjustment for dyad-level behavioral task measures, including response speed and child-parent differences in PSE, psychometric steepness, and reaction time. Child-level task behavior also showed little systematic association with the family-environment subscales. Finally, leave-one-dyad-out analyses indicated that the vmPFC- family conflict association was not driven by any single dyad. These checks do not establish a causal mechanism, but they support the interpretation that the vmPFC result reflects dyad-specific variation in affective evaluative pattern similarity rather than task performance, response quality, or general adult-child pairing.

The study also has methodological implications for research on parent-child neural similarity. Prior studies have measured neural similarity using resting-state connectome similarity, movie-evoked connectivity similarity, representational similarity during stress or empathy tasks, and real-time interbrain synchrony. These approaches capture different forms of parent-child neural alignment and differ in what they can say about shared processing. The present study uses a response-defined cross-decoding approach. It asks whether one person’s neural axis for assigning positive versus negative meaning to social-affective cues predicts the other person’s judgments. This provides a way to ask not only whether two brains are similar, but what kind of representational content is shared. This is possible because the evaluative axis is defined by within-person choice variation at matched stimulus intensity, isolating the neural signature of the judgment from that of the stimulus (Thielscher & Pessoa, 2007). Applied across dyad members, it asks whether two people share not their response to the same faces, but the way they sort faces into positive and negative. In this sense, the contribution is both substantive and methodological. Substantively, the findings point to vmPFC affective valuation as a candidate level at which perceived family conflict is reflected in parent-child neural similarity. Methodologically, they illustrate how parent-child cross-decoding can separate shared evaluative coding from broader neural similarity to the same stimuli.

### Limitations and Conclusions

Several limitations should be noted. The sample was small, and the findings should be treated as preliminary until replicated in larger samples. The design was cross-sectional and cannot establish whether family conflict leads to lower vmPFC evaluative similarity, whether divergent affective evaluation contributes to conflict, or whether both reflect other family or child-level factors. The primary family-environment effect was based on child report, so future studies should include multi-informant models, observational measures of family interaction, and longitudinal designs. The vmPFC cluster was identified by the family conflict covariate analysis, so follow-up analyses extracted from this cluster should be interpreted as descriptive characterization rather than independent confirmation. Finally, because the task used an angry-happy continuum, the current data cannot fully separate a threat-specific interpretation from a broader negative-positive valuation account.

Within these limits, the present findings provide initial evidence that youth-perceived family conflict is associated with lower neural similarity during affective judgment. The result was specific to true dyads, was not detected in face-selective perceptual regions, was not accounted for by dyad-level task behavior, and was not driven by any single dyad. These findings suggest that affective valuation, rather than sensory encoding alone, may be a representational level at which family environment is reflected in parent-child neural similarity. If replicated, these findings may have practical relevance. They locate one correlate of family conflict not in whether children perceive faces accurately, but in how parents and children converge in assigning affective meaning, which points to evaluative interpretation rather than perception as a potential focus for understanding how family conflict relates to youth affective functioning. Given the preliminary and cross-sectional nature of the evidence, this implication is offered as a direction for future work rather than a basis for intervention. More broadly, this approach shows where in the brain, and under what relational conditions, parent-child similarity carries developmental meaning.

## Supplementary Materials

### Youth’s Task Behavior and Family-Environment Subscales

To evaluate whether family environment was associated with task behavior, we correlated youth-level behavioral measures with each family-environment subscale. Youth PSE was not associated with Family Conflict (r = 0.007, p = .97), Family Cohesion (r = 0.171, p = .41), or Family Identity (r = 0.280, p = .18). Psychometric steepness was also unrelated to Family Conflict (r = 0.176, p = .40), Family Cohesion (r = −0.113, p = .59), or Family Identity (r = 0.138, p = .51). Youth mean reaction time showed a modest negative association with Family Conflict (r = −0.395, p = .050; 95% CI [−0.64, −0.13]), but was not significantly associated with Family Cohesion (r = 0.349, p = .088; 95% CI [-0.168 0.715]) or Family Identity (r = −0.067, p = .75). Thus, family environment did not systematically track response criterion or sensitivity to expression-related visual feature changes. The only behavioral association with family environment was a modest relation between higher Family Conflict and faster youth reaction time.

**Table S1.** Associations between youth task behavior and family-environment subscales.

|  | Family Conflict | Family Cohesion | Family Identity |
| --- | --- | --- | --- |
| **PSE** | $r = 0.007$, $p = .97$ | $r = 0.171$, $p = .41$ | $r = 0.280$, $p = .18$ |
| **Psychometric steepness, k** | $r = 0.176$, $p = .40$ | $r = -0.113$, $p = .59$ | $r = 0.138$, $p = .51$ |
| **Mean RT** | $r = -0.398$, $p = .049$;<br>95% CI $[-0.64, -0.13]$ | $r = 0.349$, $p = .088$;<br>95% CI $[-0.20, 0.72]$ | $r = -0.065$, $p = .76$ |

**Note.** PSE = point of subjective equality. Higher PSE indicates a more negative response criterion. Psychometric steepness, k, indexes sensitivity to expression-related visual feature changes along the morph continuum. Mean RT reflects youth mean reaction time across task trials. Pearson correlations are reported. The bootstrap confidence interval is shown for the reaction-time association with Family Conflict.

### Directional Decomposition of vmPFC Parent-Child Neural Similarity

To characterize whether the neural similarity-family conflict association in vmPFC was driven by one decoding direction, we decomposed the direction-averaged neural similarity index into child-axis to parent-trial decoding and parent-axis to child-trial decoding. Both directional components showed negative associations with family conflict. Child-axis to parent-trial decoding was negatively associated with family conflict (r = −0.573, p = .003; 95% CI [−0.79, −0.20]), as was parent-axis to child-trial decoding (r = −0.540, p = .005; 95% CI [−0.78, −0.28]). Thus, the vmPFC neural similarity-family conflict association was not attributable to a single decoding direction, but was present across both directional components of the cross-decoding measure.

**Table S2.** Directional components of vmPFC parent-child neural similarity.

| Directional component | Association with family conflict |
| --- | --- |
| Child-axis to parent-trial decoding | $r = -0.573$, $p = .003$; 95% CI $[-0.79, -0.20]$ |
| Parent-axis to child-trial decoding | $r = -0.540$, $p = .005$; 95% CI $[-0.78, -0.28]$ |

**Note.** Pearson correlations are reported. Confidence intervals are bootstrap 95% confidence intervals. Directional AUC values were averaged to form the primary dyad-level parent-child neural similarity index.

### Constituent-Scale Associations with Youth Affective Distress

We also examined the five constituent scales separately for descriptive context. Parent-child neural similarity was negatively associated with negative affect (PANAS-Negative: r = −0.711, p = .0001; 95% CI [−0.88, −0.35]) and depression (r = −0.596, p = .002; 95% CI [−0.85, −0.12]). It also showed a negative association with perceived stress by the conventional p < .05 threshold, although the bootstrap confidence interval included zero (PSS: r = −0.458, p = .021; 95% CI [−0.76, 0.05]). Associations with state anxiety and trait anxiety were not significant (STAI-S: r = 0.084, p = .69; STAI-T: r = −0.325, p = .11). These constituent-scale analyses are descriptive and should be interpreted cautiously given the small sample size and the preliminary nature of the composite-level association.

**Table S3.** Associations between vmPFC parent-child neural similarity and constituent youth distress.

| Youth distress measure | Association with vmPFC parent-child neural similarity |
| --- | --- |
| Negative affect | $r = -0.711$, $p = .0001$, 95% CI $[-0.88, -0.35]$ |
| Depression | $r = -0.596$, $p = .002$, 95% CI $[-0.85, -0.12]$ |
| Perceived stress | $r = -0.458$, $p = .021$, 95% CI $[-0.76, 0.05]$ |
| State anxiety | $r = 0.084$, $p = .69$ |
| Trait anxiety | $r = -0.325$, $p = .11$ |

measures.
**Note.** Pearson correlations are reported. Bootstrap confidence intervals are shown where available. The perceived-stress association reached the conventional $p < .05$ threshold, but its bootstrap confidence interval included zero. These constituent-scale analyses are descriptive and were not used as the primary test of youth affective distress.

### Single-Trial β Estimation and Parent-Child Cross-Decoding

#### Single-trial β estimation with GLMsingle

Trial-wise neural patterns were estimated separately for each participant using GLMsingle (Prince et al., 2022; **Fig. S1A**). For each voxel, trial-level responses were estimated with an LSA-style design matrix in which each of the 196 trials was entered as a separate regressor and convolved with a candidate hemodynamic response function (HRF). The HRF was not fixed a priori. Instead, GLMsingle tested a library of 20 candidate double-gamma HRFs, which varied in peak latency, peak width, and undershoot magnitude. For each voxel, the HRF that best fit the observed BOLD time series, defined by the highest variance explained, was retained. The corresponding LSA solution provided the trial-level β estimates for that voxel.

HRF selection was therefore voxel-specific, but the selected HRF was held constant across trials within a voxel. This allowed local hemodynamic differences across voxels to be modeled without treating HRF variation as trial-by-trial signal variation. We used the TYPE-B/FITHRF β estimates as the trial-level neural patterns. GLMdenoise (TYPE-C) and fractional ridge regression (TYPE-D) were not applied. The output of this stage was, for each participant, a voxel × trial β matrix that served as the input to all subsequent multivariate analyses.

#### Searchlight cross-decoding

Searchlight cross-decoding was conducted within the a priori medial prefrontal cortex mask to identify local mPFC regions where positive versus negative evaluative patterns generalized across parent-child dyad members (**Fig. S1B**). At each gray-matter voxel within the mPFC mask, a spherical searchlight was defined around the center voxel (radius = 10 mm), yielding a local voxel × trial β submatrix. Voxels contributing to each sphere were drawn from the mPFC mask.

Within each sphere, trial-level β patterns were first mean-centered within each morph-intensity condition to reduce stimulus-driven pattern structure associated with facial-expression intensity. Specifically, for each participant and each of the seven morph levels, the average multivoxel pattern for that morph level was subtracted from each trial at the same morph level. This step reduced morph-intensity-related pattern structure and allowed the analysis to focus on trial-to-trial variation associated with whether the face was judged as positive or negative.

We then defined a participant-specific evaluative choice axis as the voxel-wise difference between the mean condition-centered pattern for trials judged as positive and the mean condition-centered pattern for trials judged as negative. For child-axis to parent-trial decoding, the child’s evaluative choice axis was applied to the parent’s condition-centered trial patterns from the same sphere. The resulting projection scores were used to compute the area under the ROC curve (AUC), indexing whether parent positive-response trials received higher projected scores than parent negative-response trials.

The two directional AUC values were averaged to form the dyad-level neural similarity value for that sphere, with AUC = 0.5 indicating chance-level cross-decoding. The direction-averaged AUC value was assigned to the center voxel of the searchlight. Repeating this procedure across voxels within the mPFC mask produced one mPFC parent-child neural similarity map per dyad. These maps were then carried forward to second-level permutation analyses testing whether cross-decoding-based neural similarity varied as a function of family environment and youth distress.

**Fig. S1.**
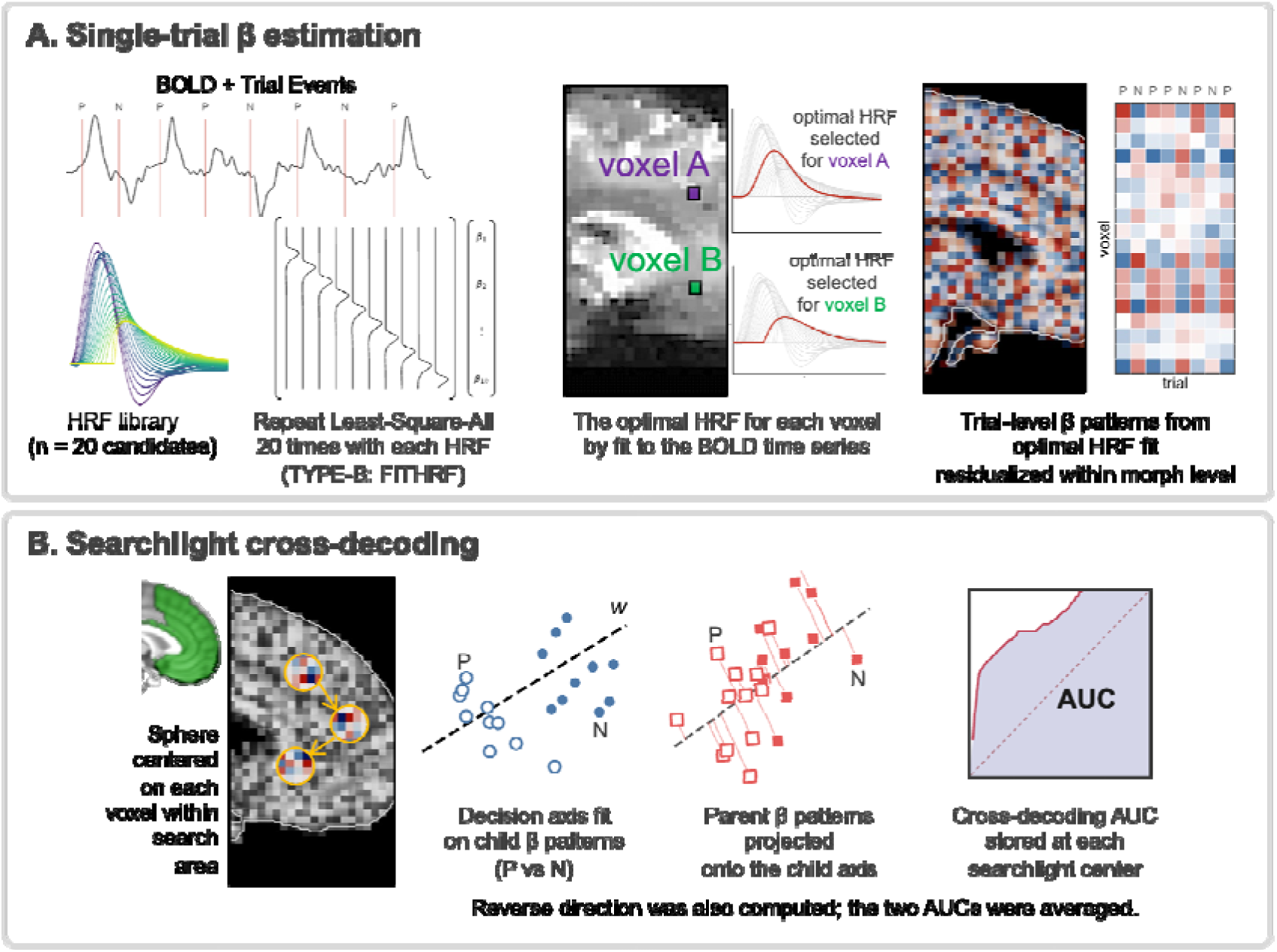
Single-trial β estimation and mPFC mask-restricted parent-child cross-decoding. **(A) Single-trial β estimation using GLMsingle TYPE-B/FITHRF.** Trial-wise neural patterns were estimated separately for each participant using GLMsingle. For each voxel, trial-level responses were estimated with an LSA-style design matrix in which each of the 196 trials was entered as a separate regressor and convolved with a candidate hemodynamic response function (HRF). The HRF was not fixed a priori. Instead, GLMsingle tested a library of 20 candidate double-gamma HRFs (Prince et al., 2022), which varied in peak latency, peak width, and undershoot magnitude. For each voxel, the HRF that best fit the observed BOLD time series, defined by the highest variance explained, was retained. The corresponding LSA solution provided the trial-level β estimates for that voxel. HRF selection was therefore voxel-specific, but the selected HRF was held constant across trials within a voxel. This allowed local hemodynamic differences across voxels to be modeled without treating HRF variation as trial-by-trial signal variation. We used the TYPE-B/FITHRF β estimates as the trial-level neural patterns. The output of this stage was, for each participant, a voxel × trial β matrix that served as the input to all subsequent multivariate analyses. **(B) Searchlight cross-decoding within the a priori mPFC mask.** Searchlight cross-decoding was conducted within the a priori medial prefrontal cortex mask to identify local mPFC regions where positive versus negative evaluative patterns generalized across parent-child dyad members. At each gray-matter voxel within the mPFC mask, a spherical searchlight was defined around the center voxel (radius = 10 mm), yielding a local voxel × trial β submatrix. Voxels contributing to each sphere were drawn from the mPFC mask. Within each sphere, trial-level β patterns were first mean-centered within each morph-intensity condition to reduce stimulus-driven pattern structure associated with facial-expression intensity. We then defined a participant-specific evaluative choice axis as the voxel-wise difference between the mean condition-centered pattern for trials judged as positive and the mean condition-centered pattern for trials judged as negative. For child-axis to parent-trial decoding, the child’s evaluative choice axis was applied to the parent’s condition-centered trial patterns from the same sphere. The resulting projection scores were used to compute the area under the ROC curve (AUC) for classifying the parent’s positive versus negative judgments. The same procedure was repeated in the opposite direction, applying the parent’s evaluative choice axis to the child’s trial patterns. The two directional AUC values were then averaged to form the dyad-level neural similarity value for that sphere, with AUC = 0.5 indicating chance-level cross-decoding. The direction-averaged AUC value was assigned to the center voxel of the searchlight. Repeating this procedure across voxels within the mPFC mask produced one mPFC parent-child neural similarity map per dyad. These maps were then carried forward to second-level permutation analyses testing whether cross-decoding-based neural similarity varied as a function of family environment and youth distress.

**Fig. S2.**
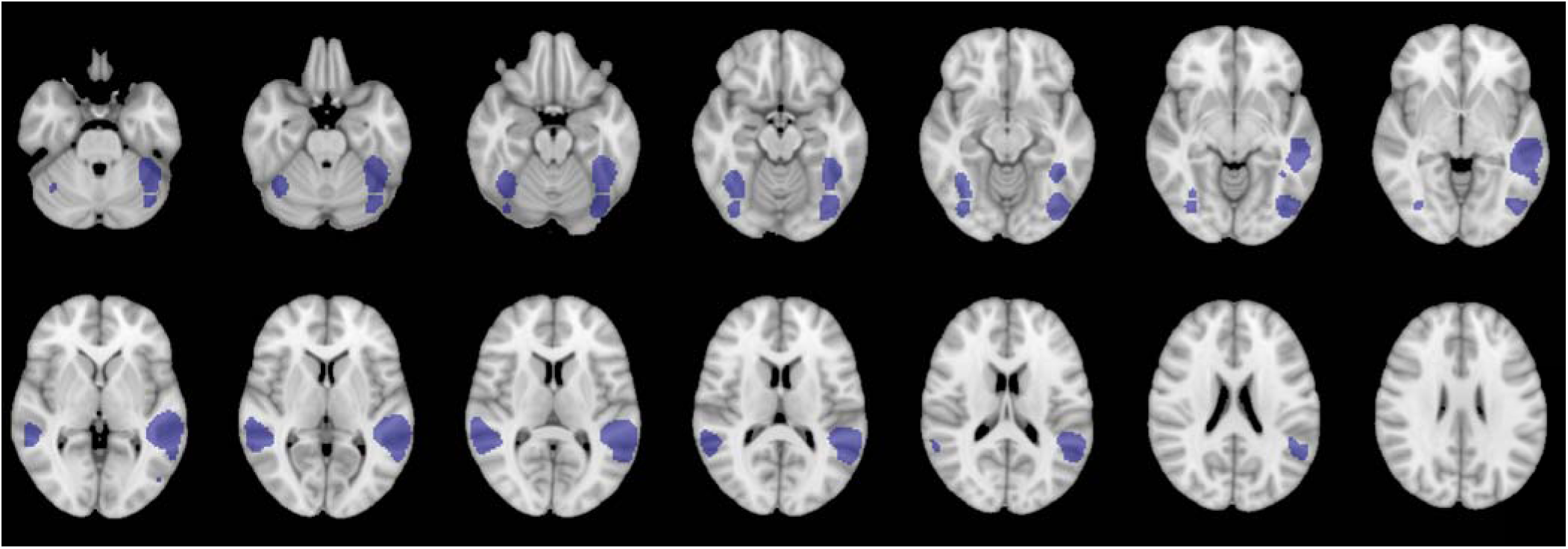
Face-selective network mask. Face-selective network parcellation from Julian, Fedorenko, Webster, and Kanwisher (2012), covering bilateral face-selective regions in occipitotemporal cortex including FFA, OFA, and posterior STS (5,900 voxels in MNI 2-mm space). Blue overlay = the mask; gray = MNI152 template.

### Leave-One-Dyad-Out Robustness Analysis

We conducted leave-one-dyad-out analyses to test whether the neural similarity - family conflict association was driven by any individual dyad. Across the 25 leave-one-dyad-out iterations, the cluster-mean correlation between parent-child neural similarity and family conflict remained negative and significant in every iteration (r range = −0.769 to −0.612; M = −0.710, SD = 0.027). The directional components were similarly stable. Child-axis to parent-trial decoding remained negatively associated with Family Conflict in all iterations (r range = −0.626 to −0.461), as did parent-axis to child-trial decoding (r range = −0.636 to −0.439). In contrast, the association between vmPFC parent-child neural similarity and the youth affective distress composite was less stable. This association remained negative in every iteration but varied more in magnitude (r range = −0.520 to −0.243; M = −0.478, SD = 0.053) and lost significance in 1 of 25 iterations. These results support the robustness of the neural similarity-Family Conflict association and are consistent with the preliminary interpretation of the youth affective distress association.

**Table S4.** Leave-one-dyad-out robustness results.

| Association | LOO $r$ range | Mean $r$ | SD | Stability |
| --- | --- | --- | --- | --- |
| Neural similarity × Family Conflict | $-0.769$<br>to $-0.612$ | $-0.710$ | 0.027 | Significant in 25/25 iterations |
| Child-axis to parent-trial decoding<br>× Family Conflict | $-0.626$<br>to $-0.461$ | $-0.573$ | 0.035 | Significant in 25/25 iterations |
| Parent-axis to child-trial decoding<br>× Family Conflict | $-0.636$<br>to $-0.439$ | $-0.540$ | 0.033 | Significant in 25/25 iterations |
| Neural similarity ×<br>youth affective distress | $-0.520$<br>to $-0.243$ | $-0.478$ | 0.053 | Negative in 25/25 iterations;<br>nonsignificant in 1/25 |

**Note.** LOO = leave-one-dyad-out. Each iteration removed one dyad and recomputed the association.

### Dyad-Level Behavioral Covariate Controls

We tested whether the vmPFC neural similarity-family conflict association was accounted for by dyad-level task-behavioral characteristics. Four behavioral covariates were examined: dyad mean reaction time, absolute parent-youth difference in point of subjective equality (PSE), absolute parent-youth difference in psychometric steepness, and absolute parent-youth difference in reaction time.

The neural similarity-family conflict association in vmPFC remained significant after controlling for each covariate separately, with partial correlations ranging from r = −0.688 to r = −0.709, all ps < .001. Of the four covariates, only dyad mean reaction time was associated with Family Conflict (r = −0.438, p = .029; 95% CI [−0.70, −0.13]), with higher Family Conflict associated with faster average responses across the dyad. The neural similarity-family conflict association in vmPFC remained significant after controlling for dyad mean reaction time (partial r = −0.688, p = .0002; 95% CI [−0.88, −0.41]). The remaining covariates were not significantly associated with Family Conflict, and controlling for each left the vmPFC neural similarity-family conflict association essentially unchanged.

**Table S5.** Dyad-level behavioral covariate controls for the vmPFC neural similarity-family conflict association.

| Behavioral covariate | Covariate association with Family Conflict | Partial association between neural similarity and Family Conflict |
| --- | --- | --- |
| Dyad mean RT | $r = -0.438$, $p = .029$;<br>95% CI [-0.70, -0.13] | partial $r = -0.688$, $p = .0002$;<br>95% CI [-0.88, -0.41] |
| Parent-youth PSE difference | $r = -0.108$, $p = .61$ | partial $r = -0.706$, $p < .001$<br>95% CI [-0.88, -0.45] |
| Parent-youth psychometric steepness difference | $r = 0.364$, $p = .073$;<br>95% CI [-0.01, 0.71] | partial $r = -0.689$, $p < .001$<br>95% CI [-0.86, -0.46] |
| Parent-youth RT difference | $r = 0.065$, $p = .76$ | partial $r = -0.709$, $p < .001$<br>95% CI [-0.88, -0.47] |

**Note.** Pearson correlations are reported for associations between each behavioral covariate and family conflict. Partial correlations indicate the association between vmPFC parent-child neural similarity and family conflict after controlling for each covariate separately. PSE = point of subjective equality. RT = reaction time.

## Author Contributions

THL developed the study concept, analyzed and interpreted the data, and drafted the manuscript. YYC, QL, BY, ZZ, and YQ critically revised the manuscript. All authors approved the final version of the manuscript for submission.

## Acknowledgements

This work was supported by a Virginia Tech Institute for Society, Culture and Environment research award. In preparing this manuscript, the authors used Claude for language editing, sentence-level revision, and assistance with code generation used to render figure layouts. All data analyses, scientific content, interpretations, and conclusions were developed by the authors, who reviewed and approved the final manuscript and take full responsibility for its content.

